## Supplemental Tables, Figures and Dataset Description for "Structure-Preserving Visualization for Single-cell RNA-Seq Profiles Using Deep Manifold Transformation with Batch-Correction"

#### Supplementary Note 1: Global/hierarchical and local structure preservation criteria

Following Poin_maps[1], to quantitatively compare the performance of different embedding methods, we use scale-independent quality criteria proposed by Lee and Verleysen[2]. The main idea is that a good dimensionality reduction method will have good preservation of local and global distances on the manifold, e.g., close neighbors should be placed close to each other while maintaining large distances between distant points. Below we provide a short summary of how to compute this metric. All the details can be found in the original paper of Poin_maps[1] and Lee et al.[2].

DV receives a scRNA-seq dataset $\mathscr{D} = \left\{(\mathbf{x_i}, \mathbf{y_i})_{i=1}^{N}\right\}$ as input, where $\mathbf{x_i} \in \mathbb{R}^d$ is the gene expression vector of cell $i$, $\mathbf{y_i}$ is a categorical variable vector specifying the batch in which $\mathbf{x_i}$ is measured, $d$ is the number of measured genes, and $N$ is the number of cells. Let $\mathbf{X^h} = \left\{\mathbf{x_1^h}, ..., \mathbf{x_N^h}\right\}$ denote the high-dimensional data matrix and $d_{ij}^h$ denote the distance from $\mathbf{x_i^h}$ to $\mathbf{x_j^h}$ in the high-dimensional space, and $\mathbf{X^l} = \left\{\mathbf{x_1^l}, ..., \mathbf{x_N^l}\right\}$ denote the low-dimensional embedding matrix and $d_{ij}^l$ denote the distance from $\mathbf{x_i^l}$ to $\mathbf{x_j^l}$ in the low-dimensional space. Assume $d_{ij}^h = d_{ji}^h$ and $d_{ij}^l = d_{ji}^l$, the distances could be used to compute high $\rho_{ij}^h$ and low $\rho_{ij}^l$ dimensional ranks between the points:

$$\rho_{ij}^h = \mid \left\{k : d_{ik}^h < d_{ij}^h \text{ or } \left(d_{ik}^h = d_{ij}^h \text{ and } 1 \leq k < j \leq N\right)\right\} \mid \tag{1}$$

$$\rho_{ij}^l = \mid \left\{k : d_{ik}^l < d_{ij}^l \text{ or } \left(d_{ik}^l = d_{ij}^l \text{ and } 1 \leq k < j \leq N\right)\right\} \mid \tag{2}$$

where $\mid A \mid$ denotes the cardinality of set $A$. According to this definition, reflexive ranks are set to zero ($\rho_{ii}^h = \rho_{ii}^l = 0$) and ranks are unique, i.e. there are no ex aequo ranks ($\rho_{ij} \neq \rho_{ik}$ for $k \neq j$). This means that non-reflexive ranks belong to $\{1, \ldots, N-1\}$.

A co-ranking matrix $\mathbf{Q} = [q_{rs}]_{1 \leq r,s \leq N-1}$ can be defined as:

$$q_{rs} = \mid \left\{(i,j) : \rho_{ij}^h = r \text{ and } \rho_{ij}^l = s\right\} \mid \tag{3}$$

where $Q$ is the joint histogram of the ranks and is actually a sum of $N$ permutation matrices of size $N-1$. It contains all the necessary information about how ranks are preserved in a given low-dimensional representation.

As was demonstrated by Lee and Verleysen[2], the co-ranking matrix $\mathbf{Q}$ is straightforward to compute, and it could be used to compute $Q_{NX}$-scale-independent quality criteria for dimensionality reduction for a given value of $K = 1, \ldots, N-1$.

$$Q_{NX}(K) = \frac{1}{KN} \sum_{(r,s) \in \mathbb{UL}_K} q_{rs} \tag{4}$$

where $\mathbb{UL}_K = \{1, \ldots, K\} \times \{1, \ldots, K\}$ is the upper left corner of co-ranking matrix. $Q_{NX}(K) \in [0,1]$ assesses the overall quality of the embedding. Essentially, it measures the preservation of $K$-ary neighborhoods. A perfect embedding has $Q_{NX}(K) = 1$ for every $K = 1, \ldots, N-1$.

The left part of the $Q_{NX}$ curve reflects how local properties are preserved, and the right part corresponds to the preservation of global properties. To improve its readability, Lee et al. proposed to use two scalar quality criteria $Q_{local}$ and $Q_{global}$ focusing separately on low and high dimensional qualities of the embedding:

$$Q_{\text{local}} = \frac{1}{K_{\max}} \sum_{K=1}^{K_{\max}} Q_{NX}(K) \tag{5}$$

$$Q_{\text{global}} = \frac{1}{N - K_{\max}} \sum_{K=K_{\max}}^{N-1} Q_{NX}(K) \tag{6}$$

where $K_{max}$ defines the split of the $Q_{NX}(K)$ curve and is automatically computed as:

$$K_{\max} = \arg\max_K \left(Q_{NX}(K) - \frac{K}{N-1}\right) \tag{7}$$

The quantities of $Q_{local}$ and $Q_{global}$ range from 0 (bad) to 1 (good).

In this work, we adopt Euclidean distance to estimate distances $d_{ij}^h$ in the high-dimensional space (input data). For Poincaré maps, we use geodesic distances estimated as the length of a shortest-path in a $k$-nearest neighbors graph according to the original tutorial. For the distances $d_{ij}^l$ in the low-dimensional space, we use Euclidean distances for all the embeddings except DV_Poin, DV_Lor and Poincaré maps, for which we use hyperbolic distances.

#### Supplementary Note 2: Benchmarks on datasets

The cord blood mononuclear cell dataset[3] consists of 8617 cells, including 8009 cord blood mononuclear cells and 608 mouse 3T3 fibroblasts, produced by the CITE-seq protocol[4] on the 10× Chromium (v2) platform[5]. We adopt the preprocessed data in scPhere[6], which including CD14+ erythroid cells and the first ten erythrocytes in the dataset, and they selected 2000 highly variable genes based on the Seurat5 tutorial[7].

Human splenic NK cells are from a study profiling human and mouse splenic, and blood NK cells[8], and profiles by 10× Chromium (v2). We use the 1755 human splenic NK cells from donor one and adopt the preprocessed data in scPhere[6], which selected 2724 highly variable genes, partitioned the 1755 cells into four groups, and labeled them as hNK_Sp1, hNK_Sp2, hNK_Sp3, hNK_Sp4.

Human lung cells are from asthma patients and healthy controls[9], and profiles by either 10× Chromium or Drop-seq[10]. We use the preprocessed 3314 cells in scPhere[6] from a donor prepared by the Drop-seq protocol.

The mouse white adipose tissue stromal cell dataset contains 1378 cells from mouse white adipose tissue[11] profiled by 10× Chromium (v2). In the original study, the authors only analyzed 1045 tdTomato- mGFP+ cells and identified adipocyte precursor cells (APC), fibro-inflammatory progenitors (FIP), committed preadipocytes, and mesothelial cells. We adopt the preprocessed data in scPhere[6], which analyzed all the cells and further identified pericytes, macrophages, and two groups of doublets.

The reginal ganglion cell atlas (RGC) dataset consists of 35,699 mouse retinal ganglion cells profiled by 10× Chromium (v2)[12]. The original analysis identified 45 clusters, and one cluster consisted of two cell types. We adopt the preprocessed data in scPhere[6], which selected 2724 highly variable genes.

We use 599,926 human cell landscape cells (HCL)[13] from human fetal or adult tissues profiles by the Microwell-Seq platform[14]. In the original study, these cells were portioned into 102 clusters, and 77 of 102-cell clusters can be grouped into six major cell groups: fetal stromal cells, fetal epithelial cells, adult endothelial cells, endothelial cells, adult stromal cells and immune cells. We adopt the preprocessed data in scPhere[6], which selected 2724 highly variable genes. Specially, For heterogeneous case in "batch invariant" model verification experiment, we input the raw data due to the demand of whole gene names. We reclassify the training set and test set according to tissue name to satisfy complex multilevel batch factors. The 43 tissues in training set include 'AdultAdrenalGland_3', 'AdultBladder_2', 'AdultEsophagus_2', 'AdultGallbladder_1', 'AdultHeart_2', 'AdultKidney_2', 'AdultKidney_3', 'AdultLiver_2', 'AdultLiver_4', 'AdultLung_1', 'AdultLung_3', 'AdultOmentum_1', 'AdultOmentum_2', 'AdultPeripheralBlood_2', 'AdultPeripheralBlood_4', 'AdultStomach_2', 'AdultStomach_3', 'AdultThyroid_2', 'BoneMarrow_2', 'CordBloodCD34P_2', 'CordBlood_2', 'FetalAdrenalGland_2', 'FetalAdrenalGland_3', 'FetalBrain_3', 'FetalBrain_4', 'FetalBrain_5', 'FetalHeart_1', 'FetalIntestine_1', 'FetalIntestine_4', 'FetalIntestine_5', 'FetalIntetsine_3', 'FetalKidney_3', 'FetalKidney_4', 'FetalKidney_5', 'FetalLung_2', 'FetalMaleGonad_2', 'FetalPancreas_1', 'FetalPancreas_2', 'FetalRib_3', 'FetalSkin_2', 'FetalStomach_2', 'FetalThymus_1' and 'Liver_1', and the 28 tissues in test set include 'AdultAdrenalGland_2', 'AdultBladder_1', 'AdultEsophagus_1', 'AdultGallBladder_2', 'AdultHeart_1', 'AdultKidney_4', 'AdultLiver_1', 'AdultLung_2', 'AdultOmentum_3', 'AdultPeripheralBlood_3', 'AdultStomach_1', 'AdultThyroid_1', 'BoneMarrow_1', 'CordBloodCD34P_1', 'CordBlood_1', 'FetalAdrenalGland_4', 'FetalBrain_6', 'FetalHeart_2', 'FetalIntestine_2', 'FetalKidney_6', 'FetalLung_1', 'FetalMaleGonad_1', 'FetalPancreas_3', 'FetalRib_2', 'FetalSkin_3', 'FetalStomach_1', 'FetalThymus_2' and 'Liver_2'. In addition, the celltypes of confusion matrix in Supplementary Fig. 9c are arranged in 'M2 Macrophage', 'AT2 cell', 'Hepatocyte/Endodermal cell', 'Gastric chief cell', 'Proliferating T cell', 'Antigen presenting cell (RPS high)', 'Fetal Neuron', 'Fetal fibroblast', 'Fetal stromal cell', 'Intermediated cell', 'Ventricle cardiomyocyte', 'Intercalated cell', 'Proximal tubule progenitor', 'Sinusoidal endothelial cell', 'Fetal epithelial progenitor', 'Fetal enterocyte ', 'Loop of Henle', 'Mast cell', 'T cell', 'Fetal endocrine cell', 'Fibroblast', 'Smooth muscle cell', 'Fetal chondrocyte', 'Monocyte', 'Dendritic cell', 'Fetal neuron', 'B cell', 'Epithelial cell (intermediated)', 'Stratified epithelial cell', 'Thyroid follicular cell', 'Ureteric bud cell', 'Erythroid progenitor cell (RP high)', 'Neutrophil (RPS high)', 'Neutrophil', 'Macrophage', 'Fetal acinar cell', 'Epithelial cell', 'Fetal mesenchymal progenitor', 'B cell (Plasmocyte)', 'Stromal cell', 'Endothelial cell (APC)', 'Goblet cell', 'Endothelial cell (endothelial to mesenchymal transition)', 'Primordial germ cell', 'Fetal skeletal muscle cell', 'Erythroid cell', 'CB CD34+', 'Fasciculata cell', 'Basal cell', 'Endothelial cell'.

We use mouse cell atlas cells (MCA)[13] profiles by the Microwell-Seq platform for heterogeneous cases in the "batch invariant" model verification experiment when training data and testing data belong to different species. We remove the batch gene background and perform analysis using the same pipeline and parameters as for the HCL described by the original tutorial.

The cells of the colon mucosa are from 68 biopsies collected from 18 ulcerative colitis patients and 12 healthy individuals[15], and profiles by 10× Chromium (either v1 or v2). We adopt the prepocessed data in scPhere[6], which obtain a total of 301,749 cells (26,678 stromal cells and glia, 64,457 epithelial cells, and 210,614 immune cells as annotated in the original study) by filtering likely low-quality cells (clusters). The cells span 12 stromal cell types/states, 12 epithelial cell types/states, and 23 immune cell types/states, identified by unsupervised clustering and manual annotations. They used Seurat to select 1307, 1361, and 1068 highly variable genes for the three major cell types, respectively. The dataset is profiled in a complex experimental design from the colon mucosa of 18 patients with ulcerative colitis (UC), a major type of inflammatory bowel disease (IBD), and 12 healthy individuals. In addition to each individual patient biopsy being a batch, there are many other factors to consider in the original study: individuals were either healthy or with UC, cells were collected separately from the epithelial and lamina propria fractions of each biopsy, and there were two replicate biopsies for each healthy individual and as a pair of inflamed and uninflamed biopsies for the UC patients (for a few UC patients, there were replicate inflamed and/or replicate uninflamed biopsies). Finally, samples were collected at two time periods, separated by over a year (analyzed as train and test data in the original study).

The C. elegans embryonic cell dataset consists of 86,024 cells[16] profiled using 10× Chromium (v2). The embryo times are partitioned into 12-time bins, and 63.5% of the cells are assigned to 36 major cell types based on annotation from GEO: GSE126954. We adopt the preprocessed data in scPhere[6] and treat the cells with embryo time in the range 100–130 as the root cells because embryo time <100 consisted mostly of germline cells that are also observed in embryo time >650.

#### Supplementary Note 3: Supplementary tables

Supplementary Tab. 1, Supplementary Tab. 2 and Supplementary Tab. 3 describe the hyperparameters used by DV_Eu, DV_Poin and DV_Lor on the experimental datasets, respectively. Moreover, Supplementary Fig. 11 verifies the robustness of DV_Eu to hyperparameter on processing large *static* scRNA-seq data with/without batch effect problem. Supplementary Fig. 12 verifies the robustness of DV_Poin and DV_Lor to hyperparameter on processing *dynamic* scRNA-seq data with batch effect problem.

**Table 1.** Hyperparameters of DV_Eu in experimental datasets.

| | Datasets | Hyperparameters | | | | |
|---|---|---|---|---|---|---|
| | | learning rate | batch size | $\nu^{vi}$ | $\gamma$ | $\beta$ |
| Single-batch | CD14+ monocytes | 0.0005 | 100 | 0.005 | 10 | |
| | Human lung cells | 0.0001 | 500 | 0.001 | 100000 | 0 |
| | Human splenic NK cells | 0.0005 | 100 | 0.001 | 1000 | 0 |
| | Mouse white adipose tissue stromal cells | 0.0005 | 100 | 0.005 | 10 | 0 |
| | Reginal ganglion cell atlas | 0.001 | 500 | 0.001 | 100000 | 0 |
| | Human cell landscape cells | 0.0005 | 2000 | 0.01 | 1000 | 0 |
| | C. elegans embryonic cells | 0.0005 | 500 | 0.001 | 10 | 0 |
| Multi-batch | Colon mucosa cells | 0.0005 | 2000 | 0.01 | 10 | 1 |
| | Stromal cells | 0.005 | 2000 | 0.05 | 100000 | 1 |
| | Epithelial cells | 0.001 | 2000 | 0.05 | 10 | 100 |
| | Immune cells | 0.001 | 500 | 0.001 | 100000 | 100 |
| | C. elegans embryonic cells | 0.005 | 3000 | 0.01 | 10 | 0.01 |
| Pre-trained DV | Stromal cells | 0.005 | 2000 | 0.01 | 100000 | 1 |
| | Epithelial cells | 0.005 | 500 | 0.005 | 10 | 10000 |
| | Immune cells | 0.001 | 500 | 0.01 | 10 | 1 |
| | Human cell landscape cells | 0.0005 | 2000 | 0.005 | 1000 | 0 |

∗ The annotations of hyperparameters can be found in *choices of hyperparameters* subsection in *Methods* section.

**Table 2.** Hyperparameters of DV_Poin in experimental datasets.

| | Datasets | Hyperparameters | | | | |
|---|---|---|---|---|---|---|
| | | learning rate | batch size | $\nu^{vi}$ | $\gamma$ | $\beta$ |
| Single-batch | CD14+ monocytes | 0.0005 | 500 | 0.01 | 10 | 0 |
| | Human lung cells | 0.005 | 100 | 0.05 | 1000 | 0 |
| | Human splenic NK cells | 0.005 | 100 | 0.001 | 10 | 0 |
| | Mouse white adipose tissue stromal cells | 0.005 | 100 | 0.005 | 10 | 0 |
| | Reginal ganglion cell atlas | 0.005 | 500 | 0.001 | 10 | 0 |
| | Human cell landscape cells | 0.005 | 1000 | 0.01 | 1000 | 0 |
| | C. elegans embryonic cells | 0.005 | 3000 | 0.001 | 100000 | 1000 |
| Multi-batch | Epithelial cells | 0.001 | 500 | 0.005 | 10 | 100 |
| | C. elegans embryonic cells | 0.0005 | 2000 | 0.001 | 10 | 10000 |

**Table 3.** Hyperparameters of DV_Lor in experimental datasets.

| | Datasets | Hyperparameters | | | | |
|---|---|---|---|---|---|---|
| | | learning rate | batch size | $v^{vi}$ | $\gamma$ | $\beta$ |
| Single-batch | CD14+ monocytes | 0.0005 | 500 | 0.005 | 10 | 0 |
| | Human lung cells | 0.005 | 100 | 0.05 | 1000 | 0 |
| | Human splenic NK cells | 0.01 | 100 | 0.001 | 10 | 0 |
| | Mouse white adipose tissue stromal cells | 0.001 | 100 | 0.005 | 10 | 0 |
| | Reginal ganglion cell atlas | 0.005 | 500 | 0.001 | 10 | 0 |
| | Human cell landscape cells | 0.001 | 1000 | 0.01 | 1000 | 0 |
| | C. elegans embryonic cells | 0.001 | 500 | 0.001 | 10 | 0 |
| Multi-batch | Epithelial cells | 0.001 | 500 | 0.005 | 10 | 100 |
| | C. elegans embryonic cells | 0.005 | 3000 | 0.001 | 1000 | 0.01 |

### Supplementary Note 4: Supplementary figures

Supplementary Fig. 1 and Supplementary Fig. 2 note the results of DV on visualizing small scRNA-seq datasets. Supplementary Fig. 3 notes the results of DV on visualizing large *static* scRNA-seq datasets labeled by cell type. Supplementary Fig. 4 notes the results of DV on addressing complex technical and biological batch for visualization and analysis in colon biopsies from healthy individuals and UC stromal patients, and Supplementary Fig. 6 notes the results from healthy individuals, UC stromal, epithelial and immune patients. Supplementary Fig. 7 and Supplementary Fig. 8 note the results of DV_Eu on using complex batch vectors to pinpoint cell types impacted by biological factors and to generate a batch-invariant reference atlas on UC stromal and epithelial datasets, respectively. Supplementary Fig. 9 generates a pre-trained reference atlas and maps heterogeneous scRNA-seq data. Supplementary Fig. 10 notes the convergence speed rapidly on large *static* and *dynamic* scRNA-seq data without batch effect problem. Supplementary Fig. 11 verifies the robustness of DV_Eu to hyperparameter on processing large *static* scRNA-seq data with/without batch effect problem. Supplementary Fig. 12 verifies the robustness of DV_Poin and DV_Lor to hyperparameter on processing *dynamic* scRNA-seq data with batch effect problem. Supplementary Fig. 13 notes the Poincaré disk embedding of DV_Poin, which highlights the progression of C. elegans embryonic cells in time without batch effect problem. Supplementary Fig. 14 and Supplementary Fig. 15 note the Poincaré disk embeddings of DV_Lor and scPhere_wn respectively, which highlight the progression of C. elegans embryonic cells in time with batch effect problem. Supplementary Fig. 16 gives the runtime of all methods when visualizing single-cell datasets (with batch effect problem) with different cell numbers. Supplementary Fig. 17 makes the ablation study of DV when visualizing single cell datasets with batch effect problem.

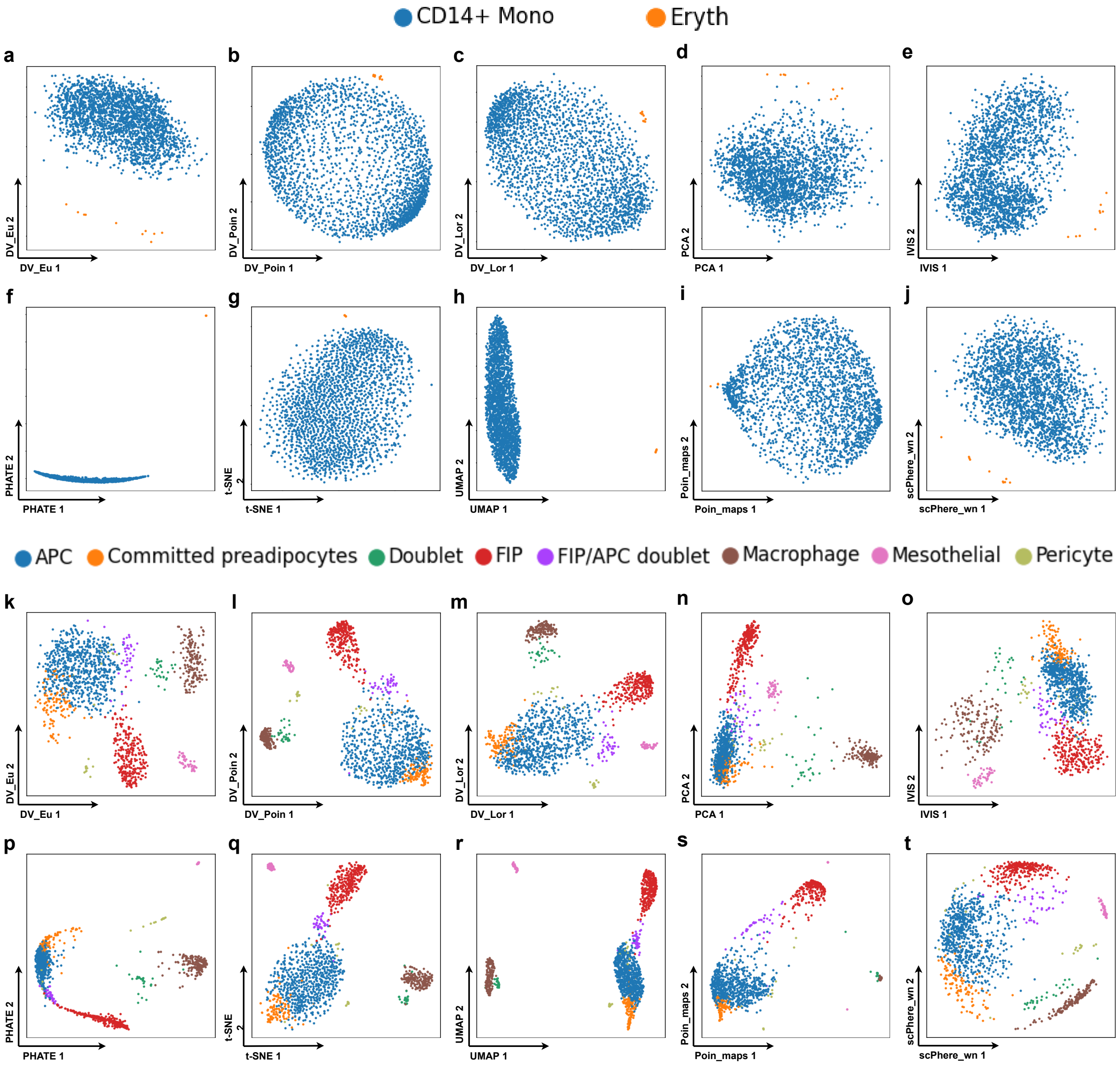


**Figure 1. DV visualizes small scRNA-seq data. a-t** DV learns latent representations that provide excellent visualization in small datasets. DV_Eu learns representations in the Euclidean space (**a, k**). DV_Poin learns representations in the hyperbolic space with Poincaré model (**b, l**). DV_Lor learns representations in the hyperbolic space with Lorentz model (**c, m**). 2-dimensional PCA (**d, n**), IVIS (**e, o**), PHATE (**f, p**), t-SNE (**g, q**), UMAP (**h, r**), Poin_maps (**i, s**) and scPhere_wn (**j, t**) for CD14+ monocytes (**a-j**) and human lung cells (GSE130148) (**k-t**) with cells colored by cell types.

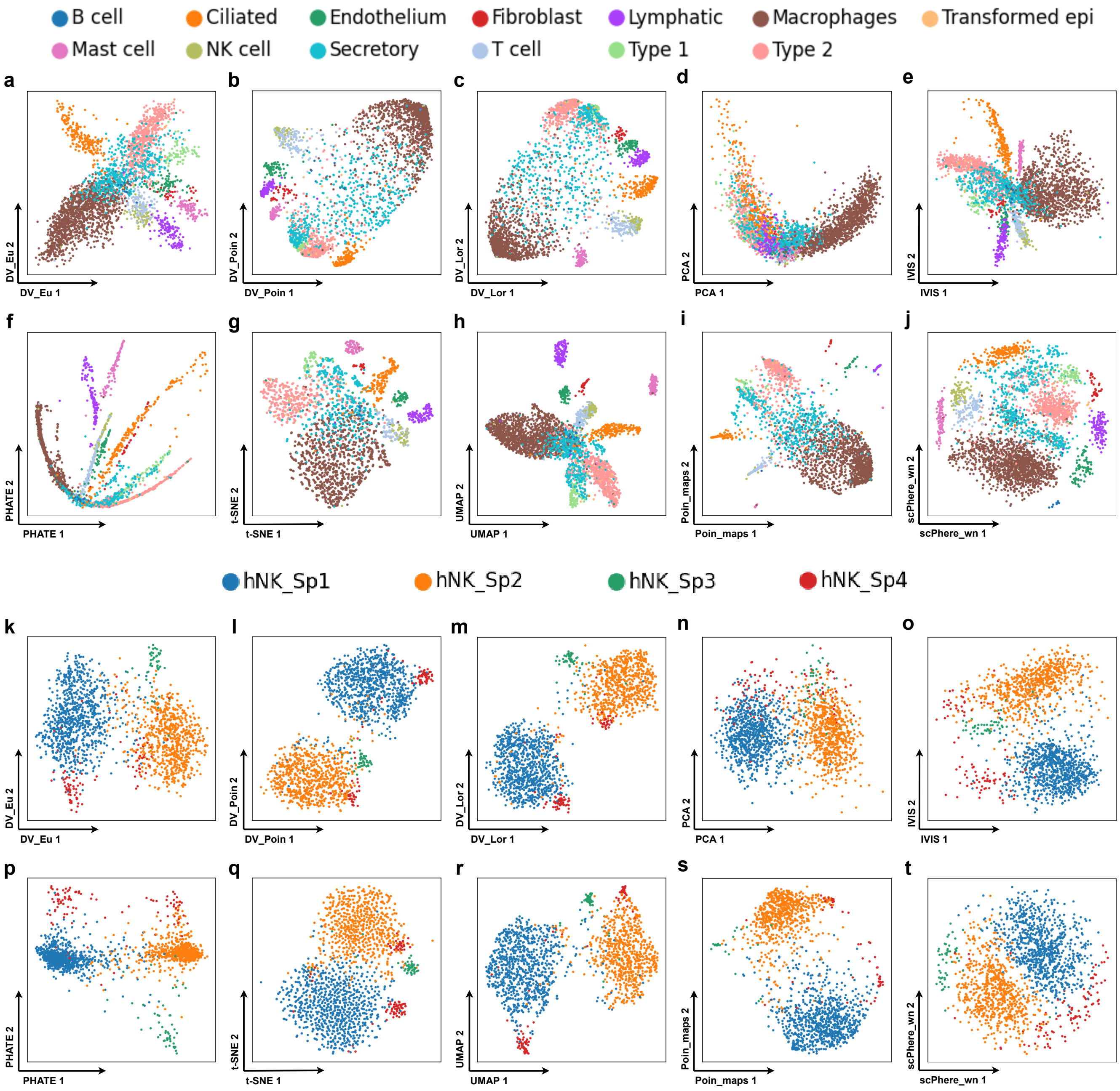


**Figure 2.** **DV visualizes small scRNA-seq data. a-t** DV learns latent representations that provide excellent visualization in small datasets. DV_Eu learns representations in the Euclidean space (**a, k**). DV_Poin learns representations in the hyperbolic space with Poincaré model (**b, l**). DV_Lor learns representations in the hyperbolic space with Lorentz model (**c, m**). 2-dimensional PCA (**d, n**), IVIS (**e, o**), PHATE (**f, p**), t-SNE (**g, q**), UMAP (**h, r**), Poin_maps (**i, s**) and scPhere_wn (**j, t**) for mouse white adipose tissue stromal cells (GSE111588) (**a-j**) and human splenic nature killer cells (GSE119562) (**k-t**) with cells colored by cell types.

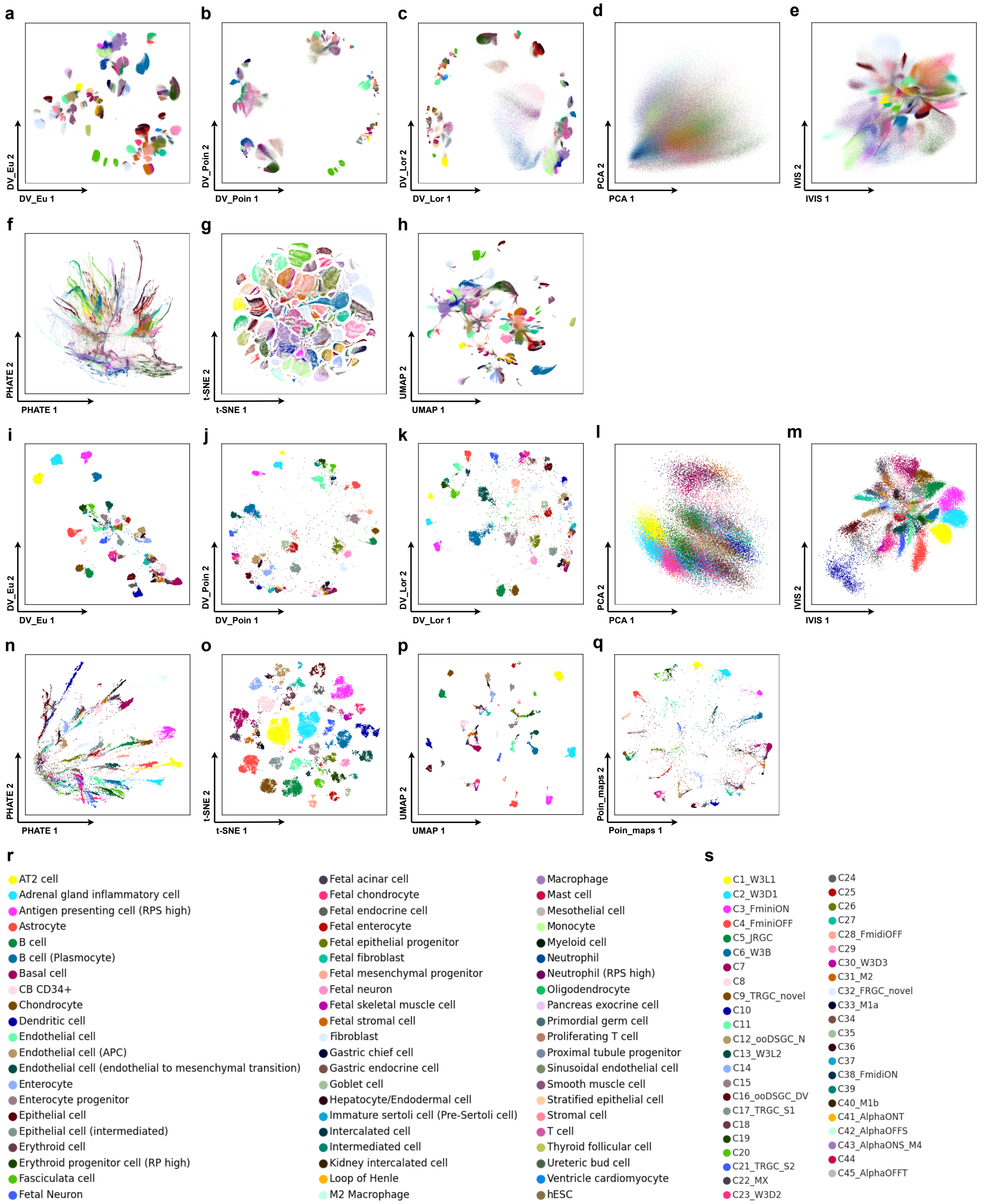


**Figure 3. DV preserves local and global structures in visualizing large *static* scRNA-seq data. a-q** DV learns latent representations that provide excellent visualization of local and global structure, even in very large datasets. DV_Eu learns representations in the Euclidean space (**a, i**). DV_Poin learns representations in the hyperbolic space with Poincaré model (**b, j**). DV_Lor learns representations in the hyperbolic space with Lorentz model (**c, k**). 2-dimensional PCA (**d, l**), IVIS (**e, m**), PHATE (**f, n**), t-SNE (**g, o**), UMAP (**h, p**) and Poin_maps (**q**) for human cell landscape (HCL) (GSE134355) (**a-h**) and 45 mouse retinal ganglion cells (RGC) (GSE137400) (**i-q**) with cells colored by cell types.

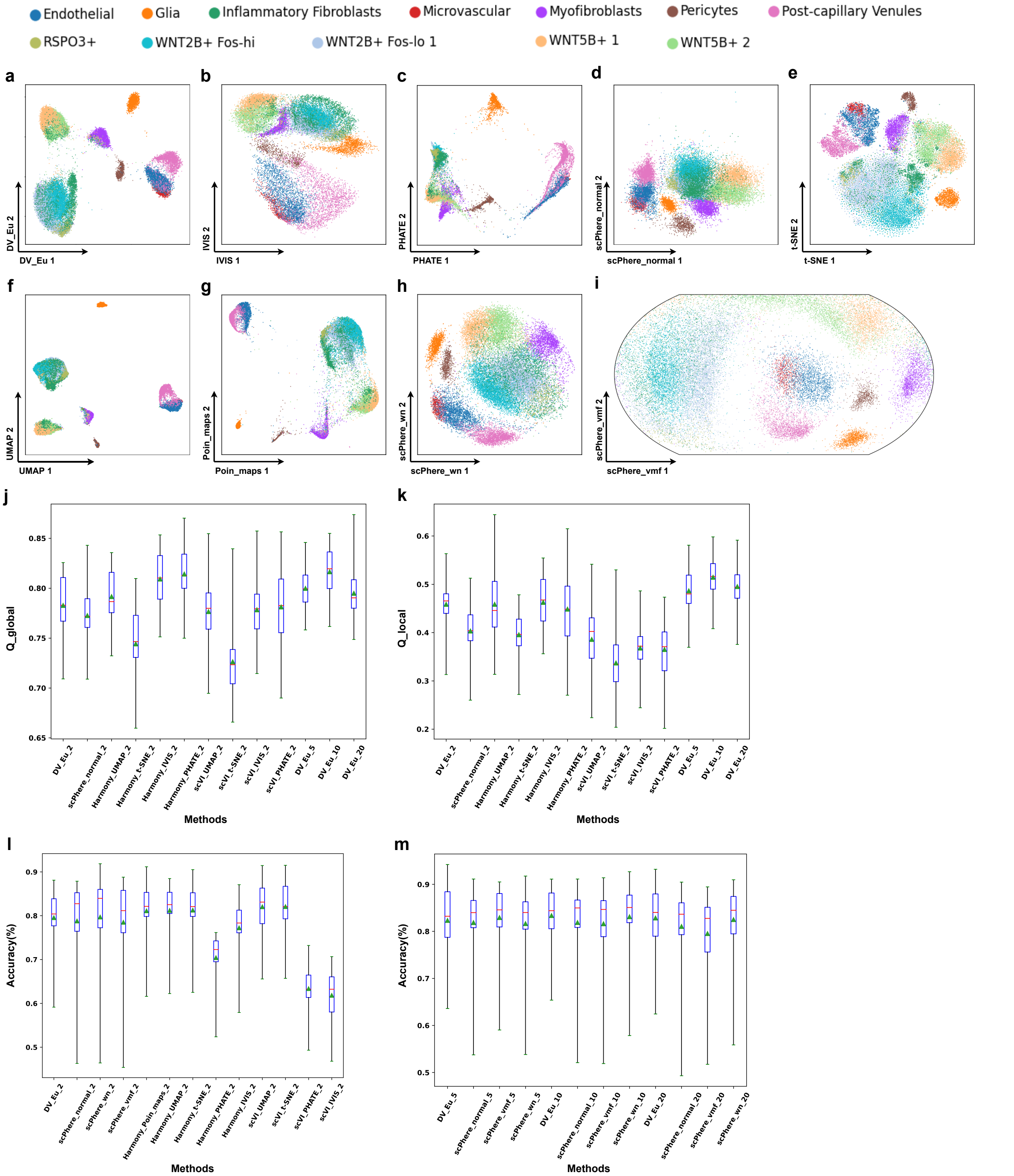


**Figure 4. DV addresses complex technical and biological batch for visualization and analysis in colon biopsies (SCP259) from healthy individuals and UC stromal patients.** 2-dimensional DV_Eu (**a**), scPhere_normal (**d**), scPhere_wn (**h**) and scPhere_vmf (**i**) embedding accounting for the patient and disease status. 2-dimensional IVIS (**b**), PHATE (**c**), t-SNE (**e**), UMAP (**f**) and Poin_maps (**g**) embedding (batch-corrected by Harmony accounting for the patient status). Successful batch correction visualization as reflected by local (**j**) and global (**k**) geometric structure preservation performance (y axis), and *k*-NN classification accuracies (y axis, *k*=5) in 2-dimensional (**l**) and low-dimensional(**m**) embeddings. The geometric structure preservation is tested on the cells from one patient between input of model and visualization embeddings. The *k*-NN classification accuracy is tested on the cells from one patient after training on the cells from all other patients. Boxplots denote means, medians and interquartile ranges (IQRs). The whiskers of a boxplot are the lowest datum still within 15 IQR of the lower quartile and the highest datum still within 15 IQR of the upper quartile.

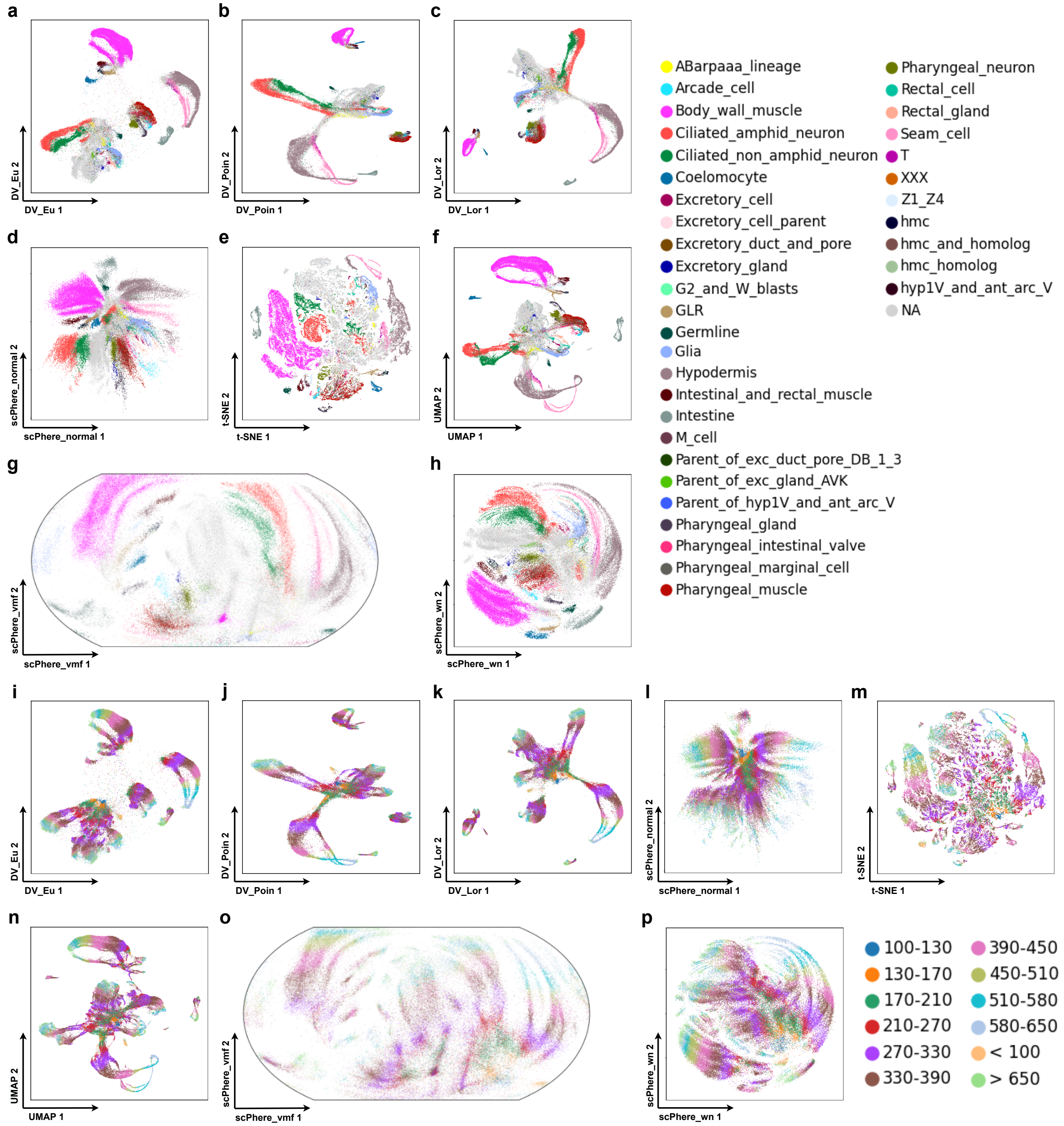


**Figure 5. DV shows developmental trajectories in visualizing large scRNA-seq differentiated data.** DV_Eu (**a, i**) and scPhere_normal (**d, l**) learn representations in the Euclidean space. DV_Poin (**b, j**) learns representations in the hyperbolic space with Poincaré model. DV_Lor (**c, k**) and scPhere_wn (**h, p**) learn representations in the hyperbolic space with Lorentz model. scPhere_vmf (**g, o**) learned representations in the hyperspherical space. 2-dimensional t-SNE (**e, m**) and UMAP (**f, n**) for ELEGAN (GSE126954) with cells colored by cell types (**a-h**) and time phases (**i-p**).

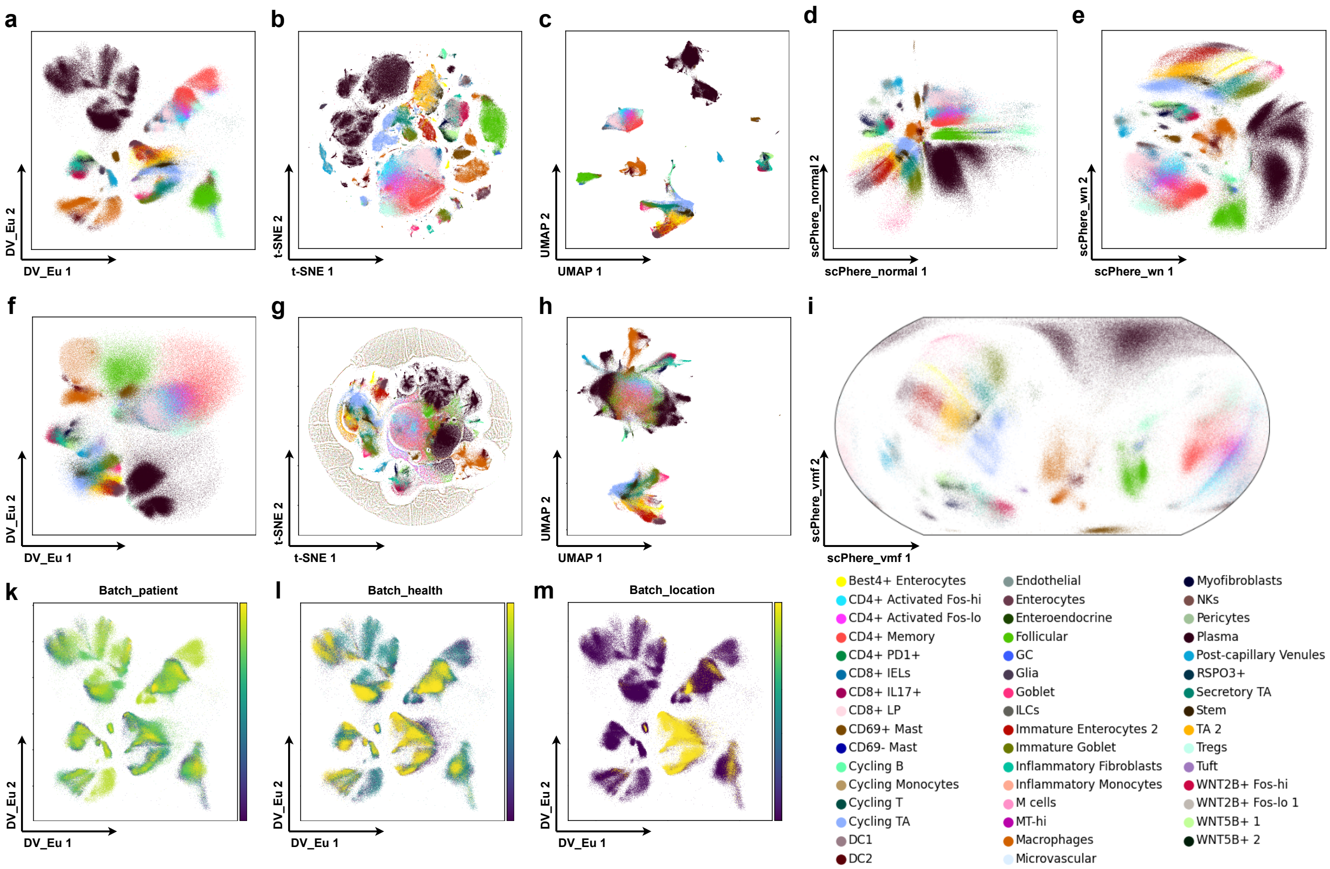


**Figure 6. DV addresses complex technical and biological batch for visualization and analysis in colon biopsies (SCP259) from healthy individuals, UC stromal, epithelial and immune patients.** DV_Eu (**a**), scPhere_normal (**d**), scPhere_wn (**e**) and scPhere_vmf (**i**) embedding accounting for the patient, location, and disease status. 2-dimensional t-SNE (**b**) and UMAP (**c**) embedding (batch-corrected by Harmony accounting for the patient status). 2-dimensional DV_Eu (**f**) (remove PCA preprocessing), t-SNE (**g**) and UMAP (**h**) embedding (batch-corrected by Harmony accounting for the patient status when removing the PCA preprocessing). DV_Eu embedding colored by batch_patient (**k**), batch_health (**l**) and batch_location (**m**) label.

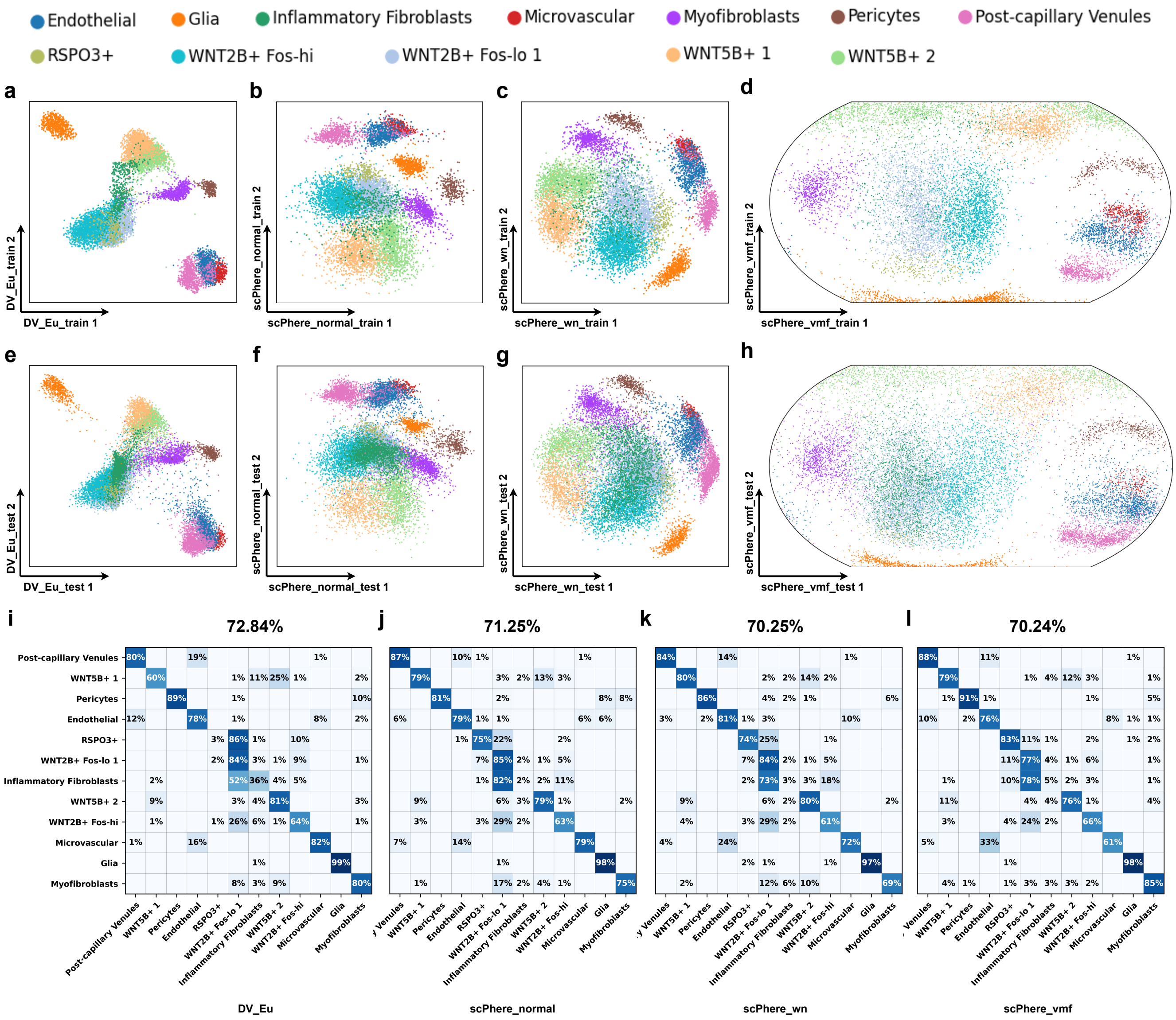


**Figure 7. Using complex batch vectors to pinpoint cell types impacted by biological factors and to generate a batch-invariant reference atlas.** Batch-invariant DV trained on 18 patients (**a-d**) and test on 12 patients (**e-h**). DV_Eu learned representations in the Euclidean space (**a, e**). 2-dimensional scPhere_normal (**b, f**), scPhere_wn (**c, g**) and scPhere_vmf (**d, h**). Confusion matrices (**i-l**) of the overlap in cells (row-centered and scaled $Z$-score, color bar) between "true" cell types from the original study (rows) and cell assignment by $k$-NN classifications (k=5) from batch-invariant DV trained only on training set for stromal cells (SCP259).

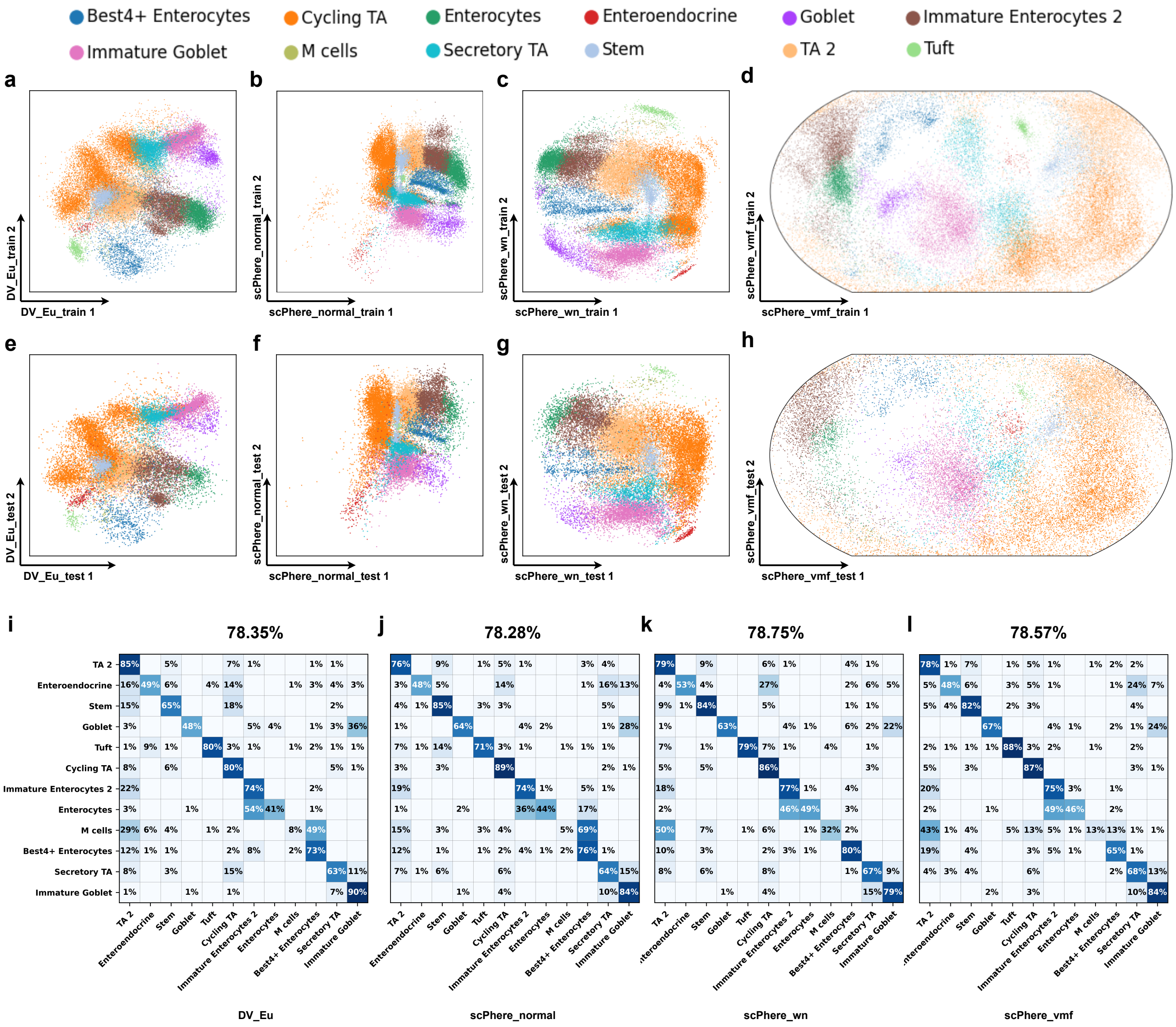


**Figure 8. Using complex batch vectors to pinpoint cell types impacted by biological factors and to generate a batch-invariant reference atlas.** Batch-invariant DV trained on 18 patients (**a-d**) and test on 12 patients (**e-h**). DV_Eu learned representations in the Euclidean space (**a, e**). 2-dimensional scPhere_normal (**b, f**), scPhere_wn (**c, g**) and scPhere_vmf (**d, h**). Confusion matrices (**i-l**) of the overlap in cells (row-centered and scaled *Z*-score, color bar) between "true" cell types from the original study (rows) and cell assignment by *k*-NN classifications (k=5) from batch-invariant DV trained only on training set for epithelial cells (SCP259).

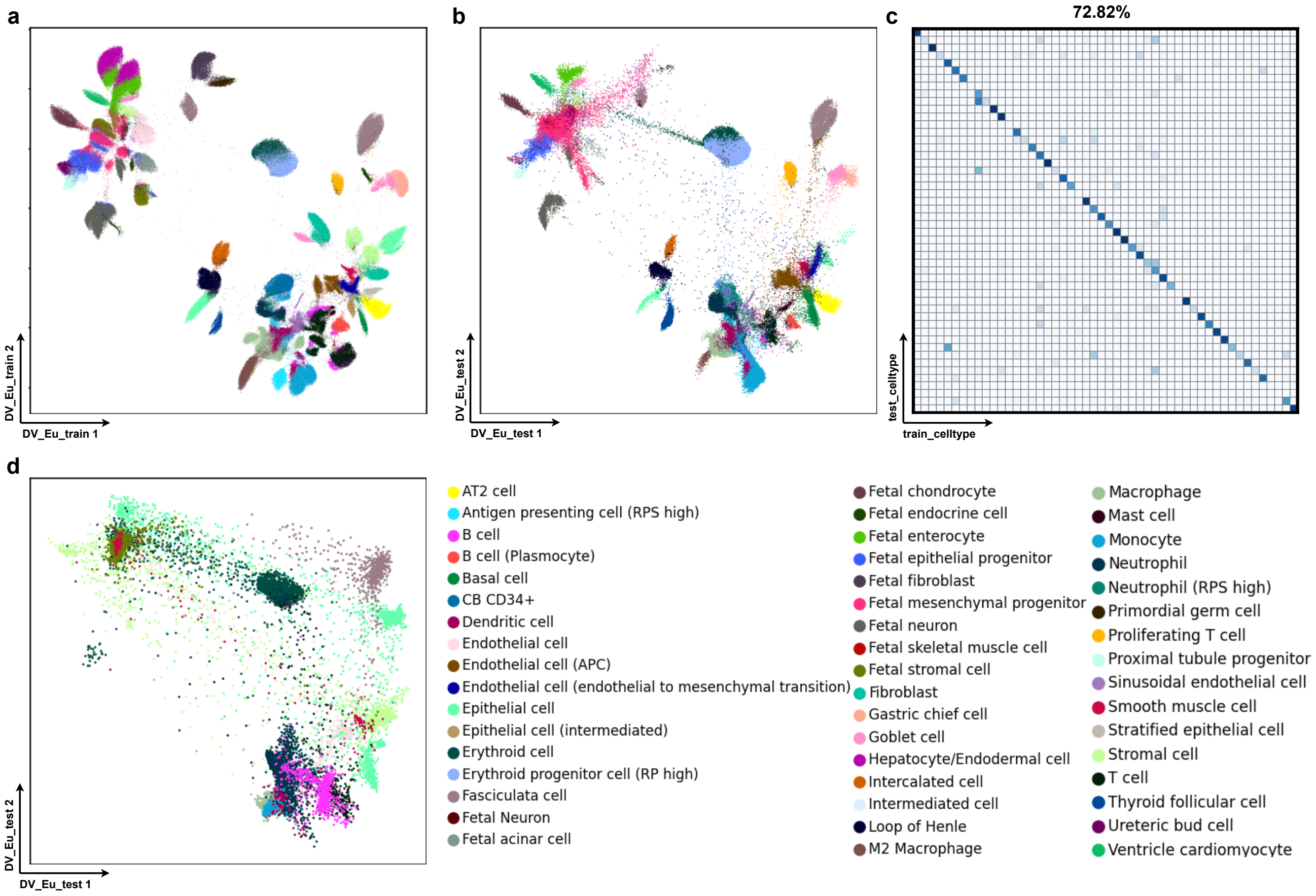


**Figure 9. DV generates a pretrained reference atlas and maps heterogeneous scRNA-seq data.** The pre-trained DV trained on HCL cells (GSE134355) from 43 tissues (**a**), and test on HCL cells from 28 tissues (**b**) and MCA cells (**d**) test data. Confusion matrices (**c**) of the overlap in cells (row-centered and scaled $Z$-score, color bar) between “true” cell types from the original study (rows) and cell assignment by $k$-NN classifications (k=5) from batch-invariant DV trained only on training set for HCL cells.

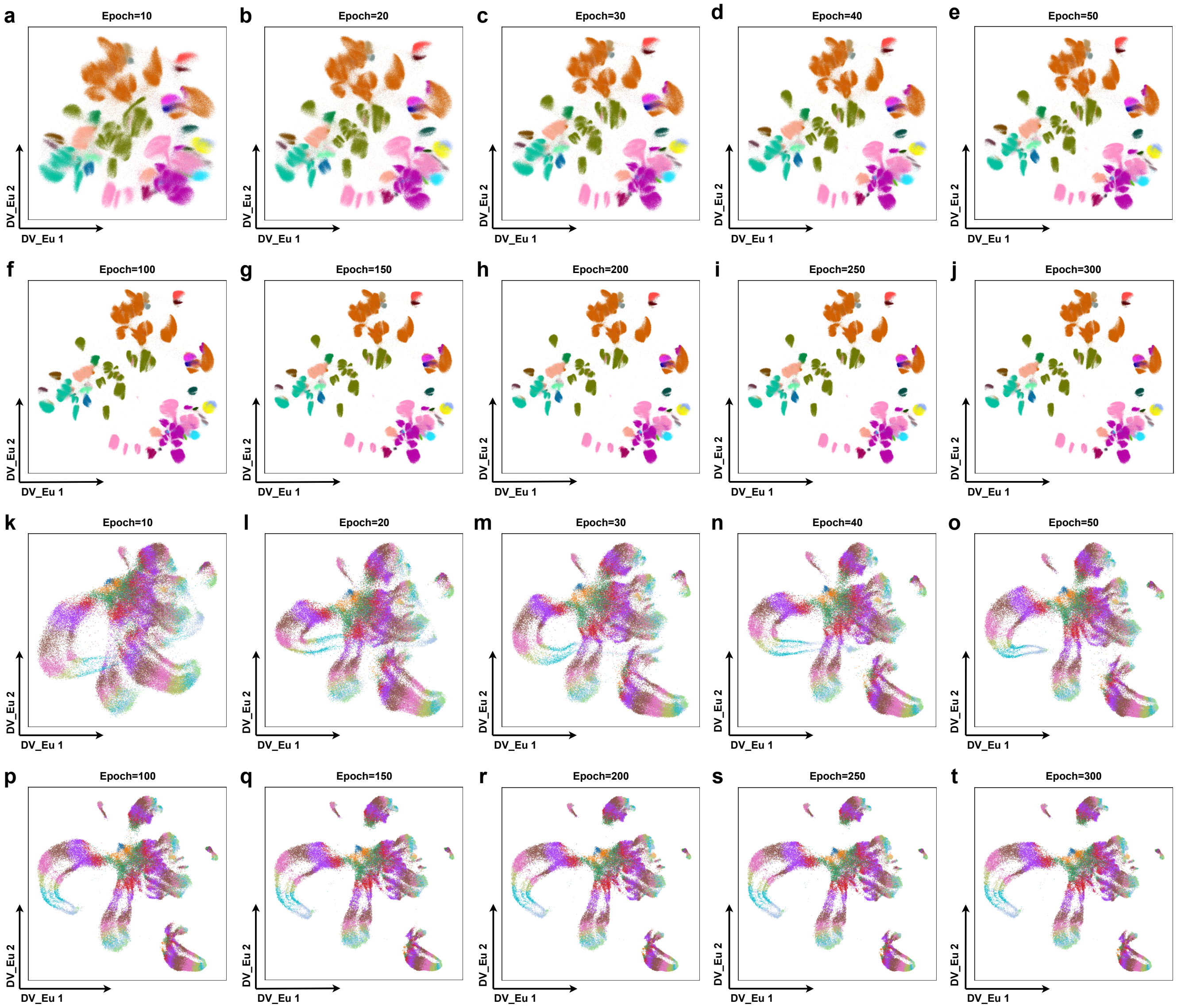


**Figure 10.** **DV converges rapidly on large *static* and *dynamic* scRNA-seq data without batch effect problem.** DV visualization results with different epochs (10, 20, 30, 40, 50, 100, 150, 200, 250, 300) for human cell landscape (HCL) (GSE134355) (**a-j**) and C. elegans embryonic (ELEGAN) (GSE126954) (**k-t**).

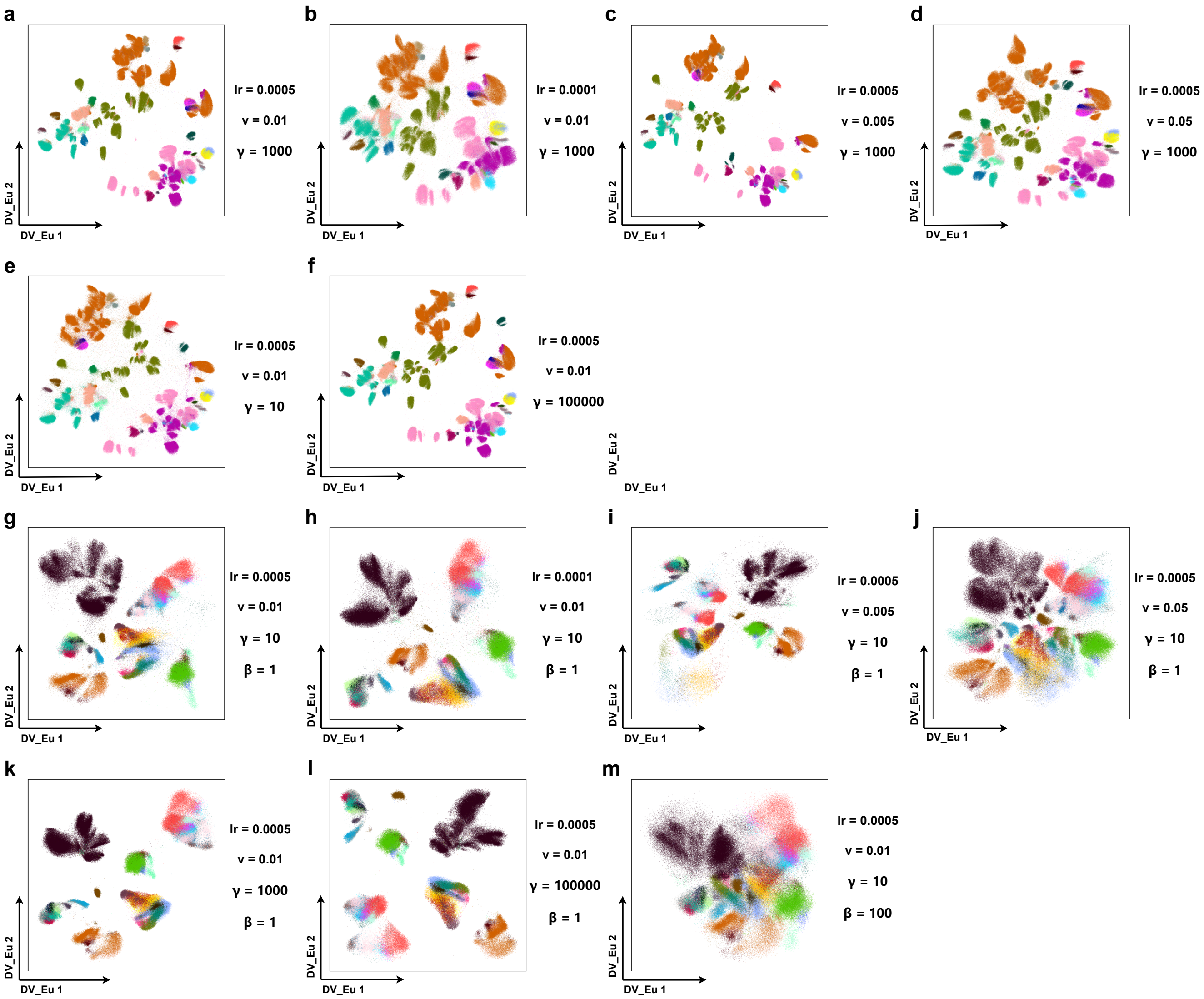


**Figure 11. DV_Eu is robust to hyperparameter on processing large *static* scRNA-seq data with/without batch effect problem.** DV visualization results with different hyperparameters (learning rate $lr$, $v$, $\gamma$, $\beta$) for human cell landscape (HCL) (GSE134355) (**a-f**) and colon biopsies (SCP259) from healthy individuals, UC stromal, epithelial and immune patients (**g-m**).

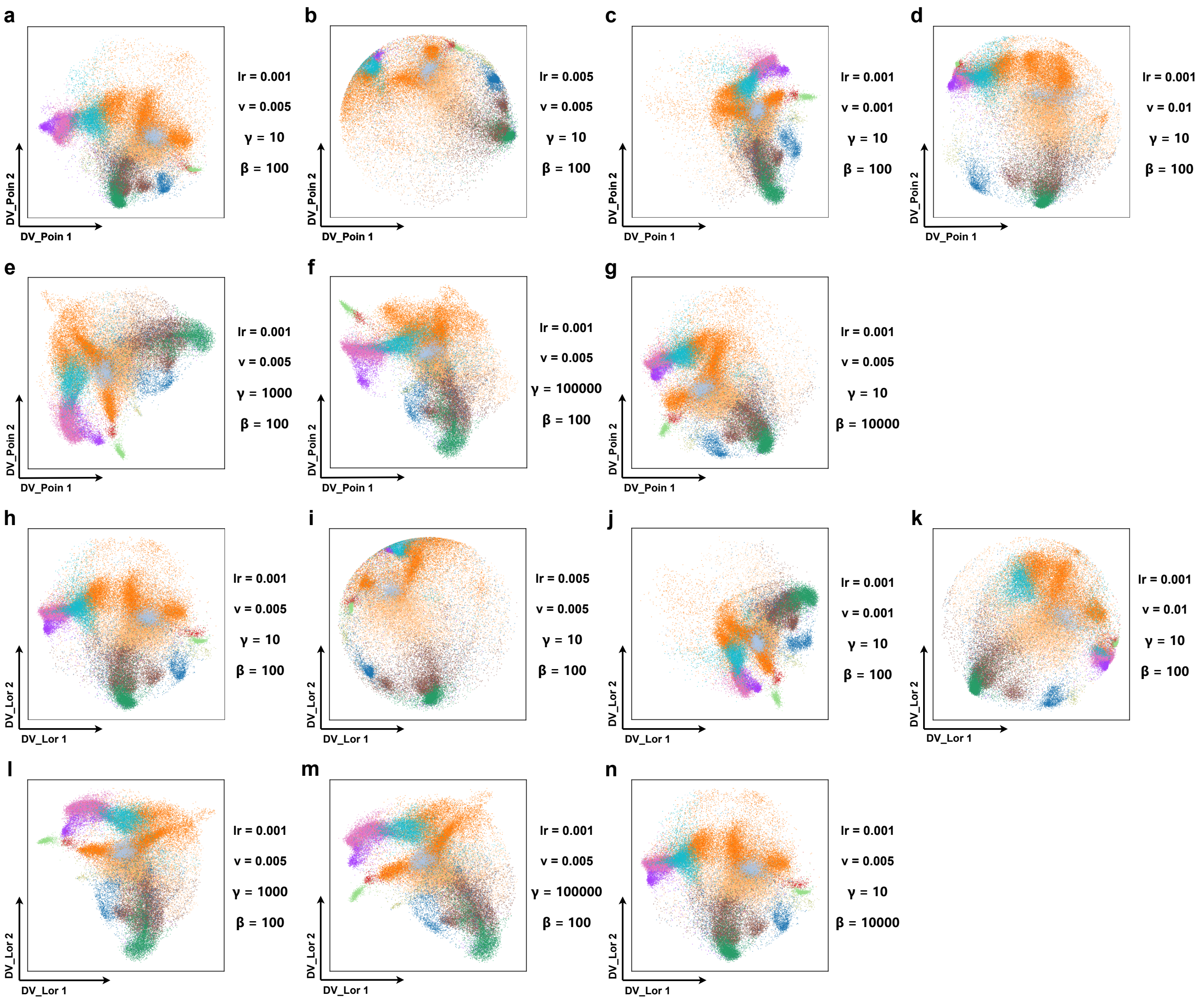


**Figure 12. DV_Poin and DV_Lor is robust to hyperparameter on processing *dynamic* scRNA-seq data with batch effect problem.** DV_Poin (**a-g**) and DV_Lor (**g-n**) visualization results with different hyperparameters (learning rate $lr$, $v$, $\gamma$, $\beta$) for colon biopsies (SCP259) from healthy individuals and UC epithelial patients.

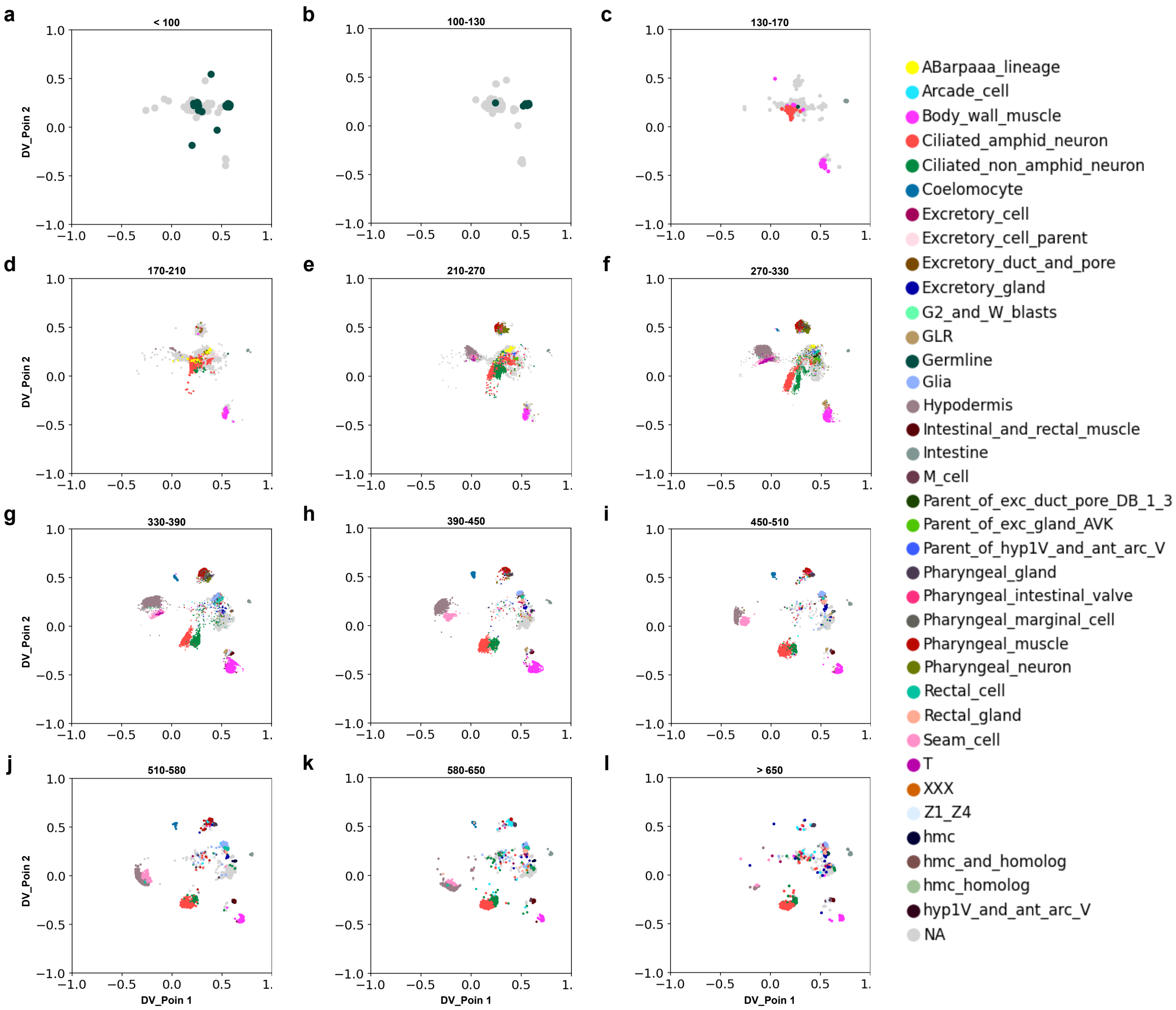


**Figure 13.** **Poincaré disk embedding of DV_Poin highlights the progression of C. elegans embryonic cells (GSE126954) in time without batch effect problem.** Embedding of all C. elegans embryonic cells in a Poincaré disk, with each panel showing only the cells from one of 12 embryonic time bins. Cells are colored by annotated cell types.

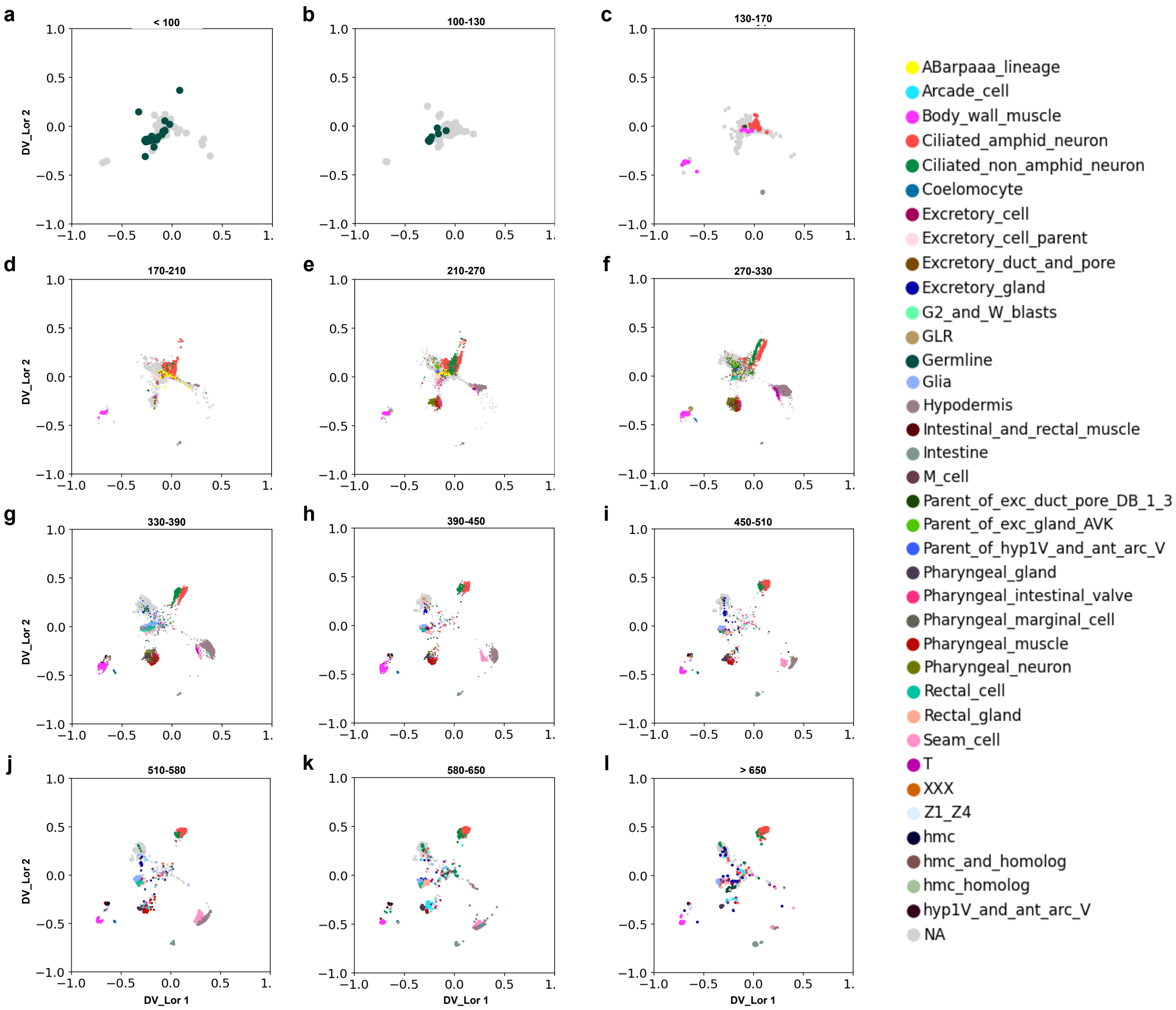


**Figure 14. Poincaré disk embedding of DV_Lor highlights the progression of C. elegans embryonic cells (GSE126954) in time with batch effect problem.** Embedding of all C. elegans embryonic cells in a Poincaré disk, with each panel showing only the cells from one of 12 embryonic time bins. Cells are colored by annotated cell types.

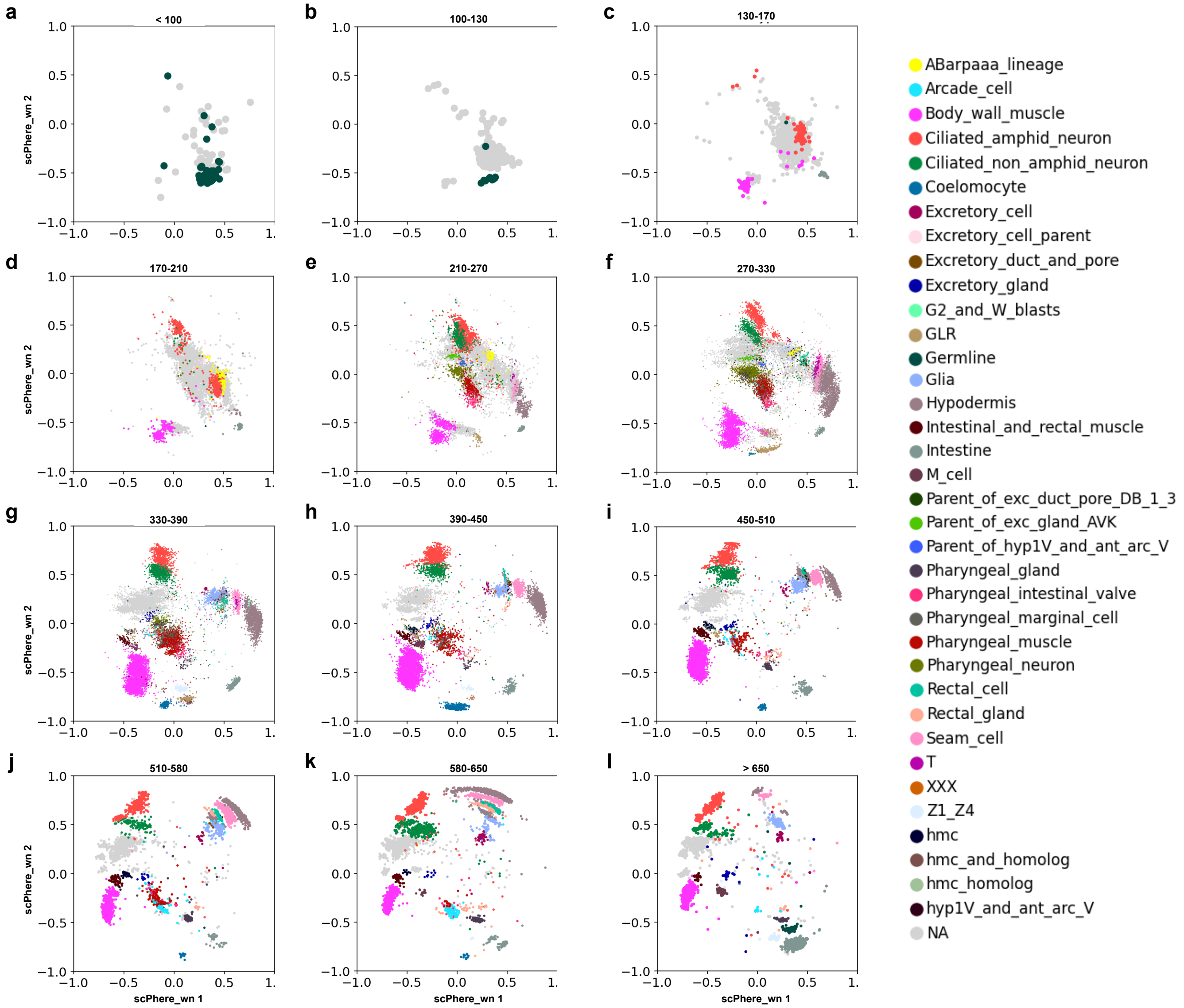


**Figure 15. Poincaré disk embedding of scPhere_wn highlights the progression of C. elegans embryonic cells (GSE126954) in time with batch effect problem.** Embedding of all C. elegans embryonic cells in a Poincaré disk, with each panel showing only the cells from one of 12 embryonic time bins. Cells are colored by annotated cell types.

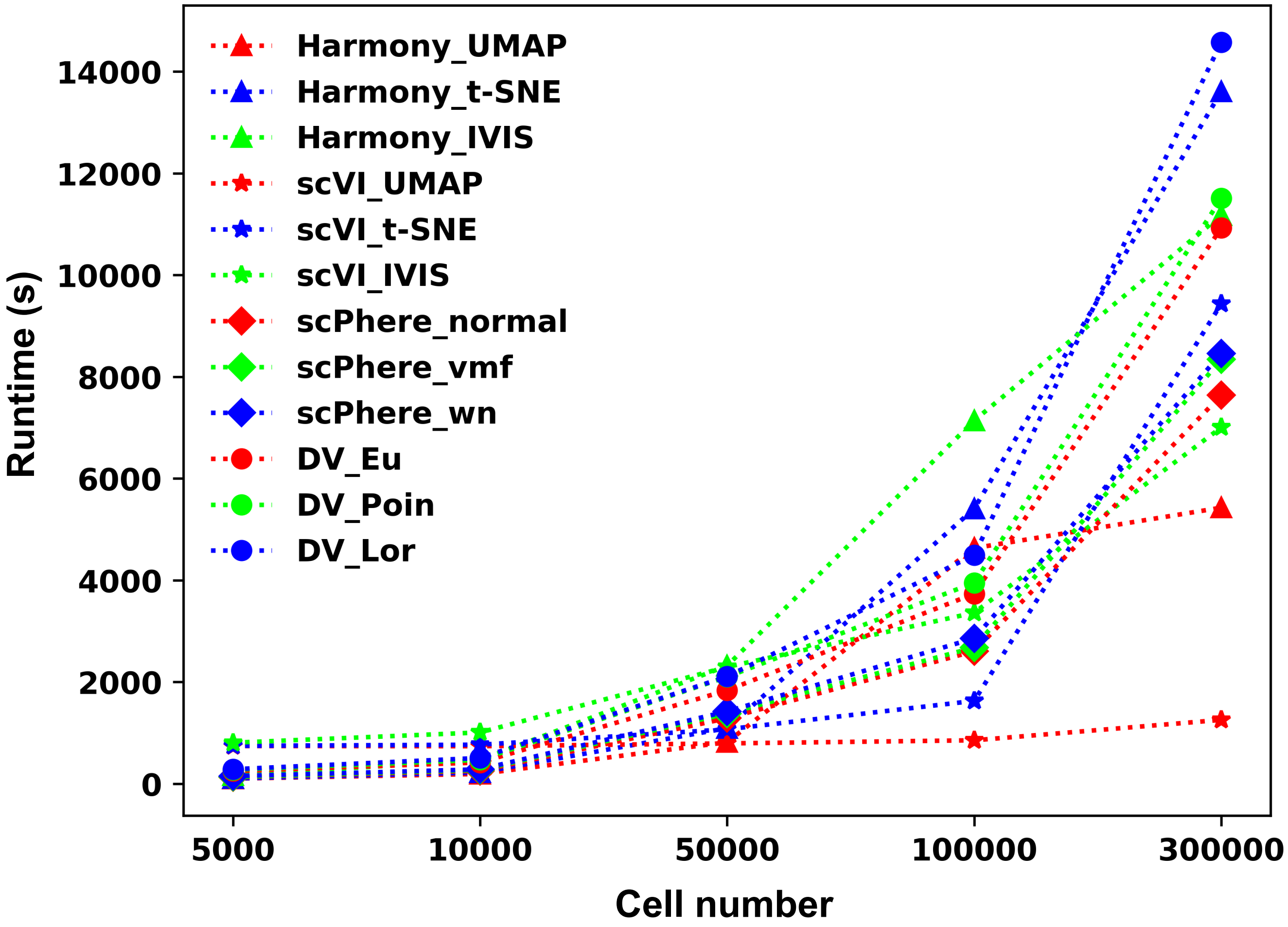


**Figure 16. Runtime of all methods when visualizing single-cell datasets (with batch effect problem) with different cell numbers.** The cell number is set to 5000, 10000, 50000, 100000 and 300000.

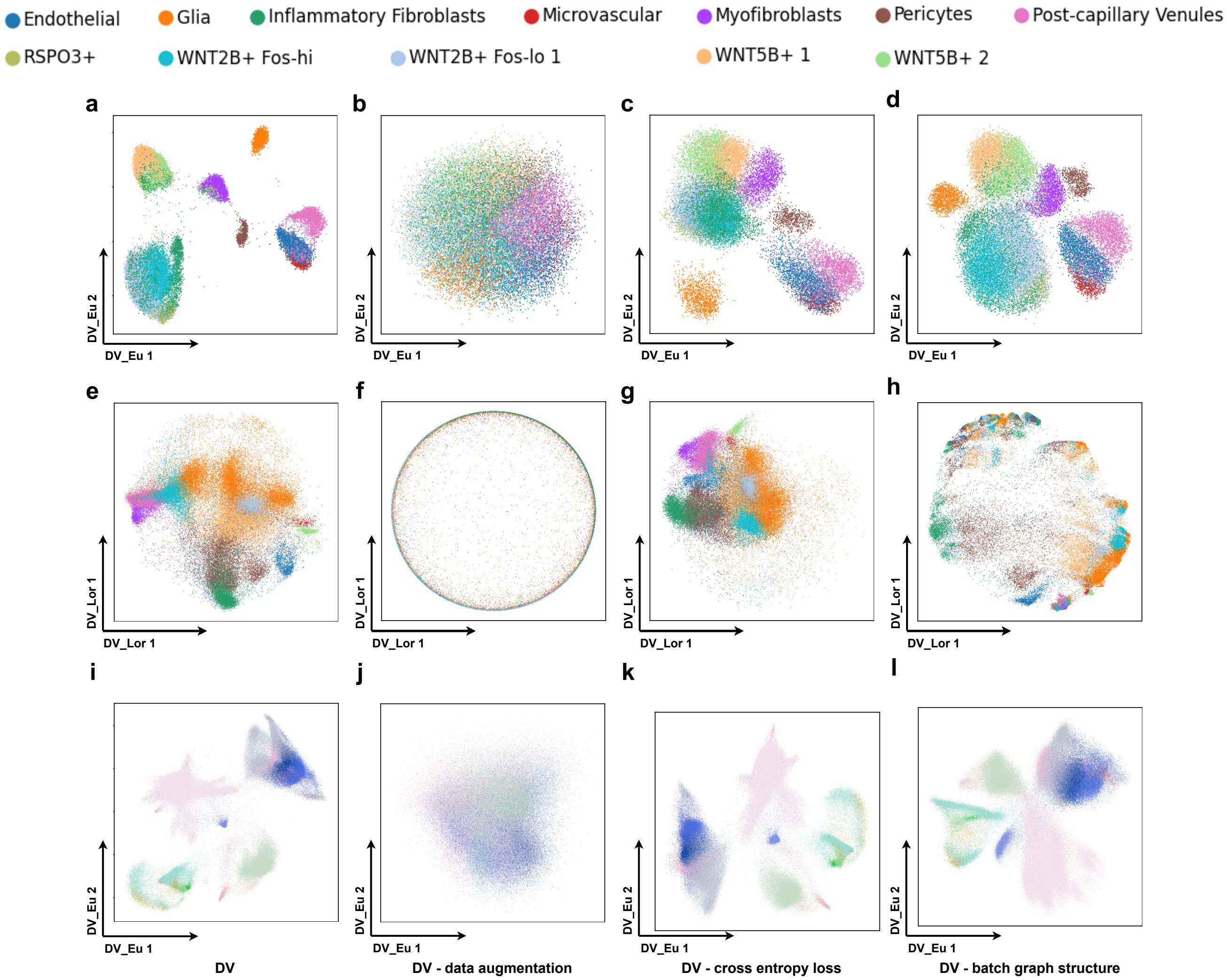


**Figure 17. Ablation study of DV when visualizing single cell datasets with batch effect problem.** We mainly assess the importance of data augmentations (**b, f, j**), cross-entropy loss (**c, g, f**) and batch graph structure (**d, h, l**) in DV_Eu on the UC stromal (**a-d**) and immune datasets (**i-l**) and DV_Lor (**e-h**) on the UC epithelial dataset.
